# A Novel Method for Calculating Beta Band Burst Durations in Parkinson’s Disease Using a Physiological Baseline

**DOI:** 10.1101/2020.03.25.008185

**Authors:** RW Anderson, RS Neuville, YM Kehnemouyi, CM Anidi, MN Petrucci, JE Parker, A Velisar, H Bronte-Stewart

**Affiliations:** Stanford University School of Medicine, Department of Neurology and Neurological Sciences, Stanford, CA, USA; Stanford University School of Medicine, Department of Neurosurgery, Stanford, CA, USA

**Keywords:** Beta fluctuations, burst durations, local field potentials, thresholding, Parkinson’s disease, Deep Brain Stimulation

## Abstract

**Background:** Pathological bursts of neural activity in Parkinson’s disease present as exaggerated subthalamic neuronal oscillations in the 8-30 Hz frequency range and are related to motor impairment.

**New Method:** This study introduces a novel method for determining burst dynamics using a baseline that matches physiological 1/f spectrum activity. We used resting state local field potentials from people with Parkinson’s disease and a simulated 1/f signal to measure beta burst durations, to demonstrate how tuning parameters (i.e., bandwidth and center frequency) affect burst durations, to compare this with high power threshold methods, and to study the effect of increasing neurostimulation intensities on burst duration.

**Results:** Burst durations calculated using the Anderson method captured the longest and broadest distribution of burst durations in a pathological beta band compared to previous methods. Mean beta band burst durations were significantly shorter on compared to off neurostimulation (p = 0.011), and their distribution was shifted towards that of the physiological 1/f spectrum during increasing intensities of stimulation.

**Comparison with Existing Method:** Existing methods of measuring local field potential power either lack temporal specificity to detect bursts (power spectral density diagrams and spectrograms) or include only higher power bursts and portions of the neural signal.

**Conclusions:** We suggest that this novel method is well suited to quantify the full range of fluctuations in beta band neural activity in the Parkinsonian brain. This method may reveal more relevant feedback biomarkers than averaged beta band power for future closed loop algorithms.

**Highlights:** A novel method for measuring variability in subthalamic local field potential oscillations in Parkinson’s disease using a physiological baseline of power.

Modeling normal brain activity using a physiological 1/f spectrum.

Burst durations depend on choice of bandwidth and center frequency.

Defining an inert frequency band whose mean burst duration overlap the physiological 1/f spectrum, from which the baseline was determined.

Burst durations progressively shortened during increasing intensities of deep brain stimulation.

## 1. Introduction

Local field potentials (LFPs) represent the summed electrical activity of local neuronal networks and have been represented classically by power spectral density (PSD) analyses and spectrograms. The PSD represents the average power of each frequency component but not the exact timing of the events that generate it. Physiological broadband neural activity is represented by a 1/f spectrum on a PSD (He, 2014; Shreve et al., 2017). Physiological neural activity consists of short duration (40 – 120 ms) neuronal oscillations and synchrony that represent normal signal processing in the sensorimotor network (Courtemanche et al., 2003; Feingold et al., 2015; Murthy and Fetz, 1996, 1992). Elevated power above the physiological 1/f spectrum within the alpha/beta band (8 – 30 Hz) represents an averaged measure of a state of higher probability of exaggerated neuronal oscillations and neural synchrony, which are pathophysiological markers of the hypokinetic state in Parkinson’s disease (PD) (Anidi et al., 2018; Bergman et al., 1994; Brown, 2003; Deffains et al., 2018; Hammond et al., 2007; Kühn et al., 2008, 2006; Velisar et al., 2019; Whitmer et al., 2012).

The classical tool used to measure how neural signals change with time is the spectrogram, which is made up of consecutive, stepped and overlapping short PSDs. Spectrograms, however, have a fixed time-frequency resolution: longer time windows result in better frequency resolution but poor temporal resolution, while short time windows allow temporal events to be easily distinguished but have poor frequency distinction. There is an emerging need to develop methods for analyzing neural signals over time that do not compromise frequency or temporal resolution.

This became pertinent for PD investigations since Feingold et al. suggested that the durations of beta band power fluctuations may be the key difference between physiological and PD related (pathological) neurophysiology (Feingold et al., 2015). Current published methods to measure the dynamics of beta band oscillatory activity have focused on event detection of a high power burst: aspects of the signal were only included in the burst analysis if and when instantaneous power rose above a threshold of the power of a bandpass filtered signal (Cagnan et al., 2019; Deffains et al., 2018; Feingold et al., 2015; Lofredi et al., 2019b; Meidahl et al., 2019; Tinkhauser et al., 2017a, 2017b, 2018; Torrecillos et al., 2018). More recently, it has been shown that burst duration, rather than burst power, was a more reliable metric to discriminate healthy from parkinsonian neural activity in non-human primates (Deffains et al., 2018).

In this study we introduce a new method (the Anderson method) to measure beta band bursts using a baseline of power. The baseline was calculated from a frequency band within the same PD LFP spectrum, which overlapped and had similar burst dynamics to the simulated physiological 1/f spectrum. This baseline, from an ‘inert’ band in the same spectrum, can be used to determine burst durations and power in any other band in the spectrum and this allows comparison of burst dynamics across subjects, as commonly used for LFP power (Anidi et al., 2018; Syrkin-Nikolau et al., 2017). We demonstrate that the Anderson method captured a broader range of burst durations (with no power discrimination) in the pathological beta band compared to other power threshold methods. The Anderson method was used to investigate the effect of increasing intensities of STN DBS on resting state beta band burst duration distributions.

## 2. Methods

### 2.1 Human subject data used to apply the Anderson method

Nine PD subjects (six male, sixteen STNs) had bilateral implantation of DBS leads (model 3389, Medtronic, Inc.) in the sensorimotor region of the STN using a standard functional frameless stereotactic technique and multipass microelectrode recording (MER) (Brontë-Stewart et al., 2010; Quinn et al., 2015). Dorsal and ventral borders of each STN were determined during MER, and the base of electrode zero was placed at the ventral border of the STN. The leads were connected through Medtronic research extensions (model 3708760) to the implanted investigative neurostimulator (Activa® PC+S, Medtronic, Inc. FDA Investigator Device Exemption approved). The preoperative selection criteria, surgical technique, and assessment of the subject have been previously described (Brontë-Stewart et al., 2010; Quinn et al., 2015). The subjects gave written informed consent to participate in the study, which was approved by the Food and Drug Administration (FDA) and the Stanford School of Medicine Institutional Review Board (IRB). Long-acting dopaminergic medication was withdrawn over 24 h (72 h for extended-release dopamine agonists), and short-acting medication was withdrawn over 12 h before the study visit. Patients had a mean age of 55.3 ± 9.5 years at the time of DBS titration experiments with an average disease duration of 8.9 ± 3.3 years. Patients had DBS surgery an average 2.4 years before experiments were conducted. Resting state data from one patient in this cohort was collected 1-year post DBS surgery.

### 2.2 Experimental protocol

Recordings were collected in the Stanford Human Motor Control and Neuromodulation laboratory. Experiments were performed off medication and the subjects were instructed to remain seated and as still as possible with their eyes open during all recordings. All resting state data was collected at least 60 minutes after DBS was turned off (Trager et al., 2016). Stimulation titration data were collected using randomized presentations of 0%, 25%, 50%, 75% and 100% of Vmax, which was the clinically equivalent DBS intensity: the intensity using a single active contact that represented the intensity being used using one or multiple contacts for clinical DBS. The last 30 seconds of a 60 second resting period at the respective stimulation intensity was used for the analysis.

### 2.3 Data acquisition and analysis

STN LFPs were recorded from electrode contact pair 0–2 (13 STNs) or 1-3 (3 STNs) on the DBS lead with LFPs high pass filtered at 0.5 Hz, low pass filtered at 100 Hz, amplified at gains set to minimize stimulation artifact and sampled at 422 Hz (10-bit resolution). Stimulation for titration experiments (60 µS pulse width, 140 Hz) were delivered through contact 1 or 2 with the maximum stimulation intensity set to the clinically equivalent single contact voltage. The power spectral density (PSD) estimate was calculated using Welch’s method with a 1-second sliding Hanning window with 50% overlap (Welch, 1967). All analysis was conducted in MATLAB (version 9.6, The MathWorks Inc. Natick, MA, USA).

### 2.4 Simulation of “physiological” neural activity

Simulation of local field potential activity in a normal brain was performed using pink (1/f) noise, generated using the MATLAB routine dsp.ColoredNoise in a period of 36 seconds, with a sample frequency of 422 samples/second and the amplitude matched to the roll off of the *in vivo* human data. Each burst method was performed on the middle 30 seconds of simulated data to eliminate filter edge effects and the simulation was repeated if any burst methods failed to find bursts in the simulated data. This sequence was repeated until 1000 episodes of simulated data with identified bursts were acquired. Filter bandwidths were tested from 1Hz-17Hz and center frequencies from 10 Hz to 24 Hz to encompass the bandwidths and center frequencies previously published in the literature.

## 3. Results

### The Anderson method – determining burst durations using a physiological baseline

#### 3.1 Establishing the physiological baseline

The goal of the Anderson method was to include all possible durations of fluctuations (bursts) in power, by determining a baseline of power, using a band in the PD spectrum, which overlapped the physiological 1/f signal. To characterize the band where the PD spectrum overlapped the simulated “normal” brain activity modeled as pink noise, we plotted both the raw LFP signal and the PSD of these signals on the same axis in Figure 1. The PD subject’s raw unfiltered LFP had elevated amplitude when compared to the pink noise data (Fig. 1A and B).

**Figure 1:**
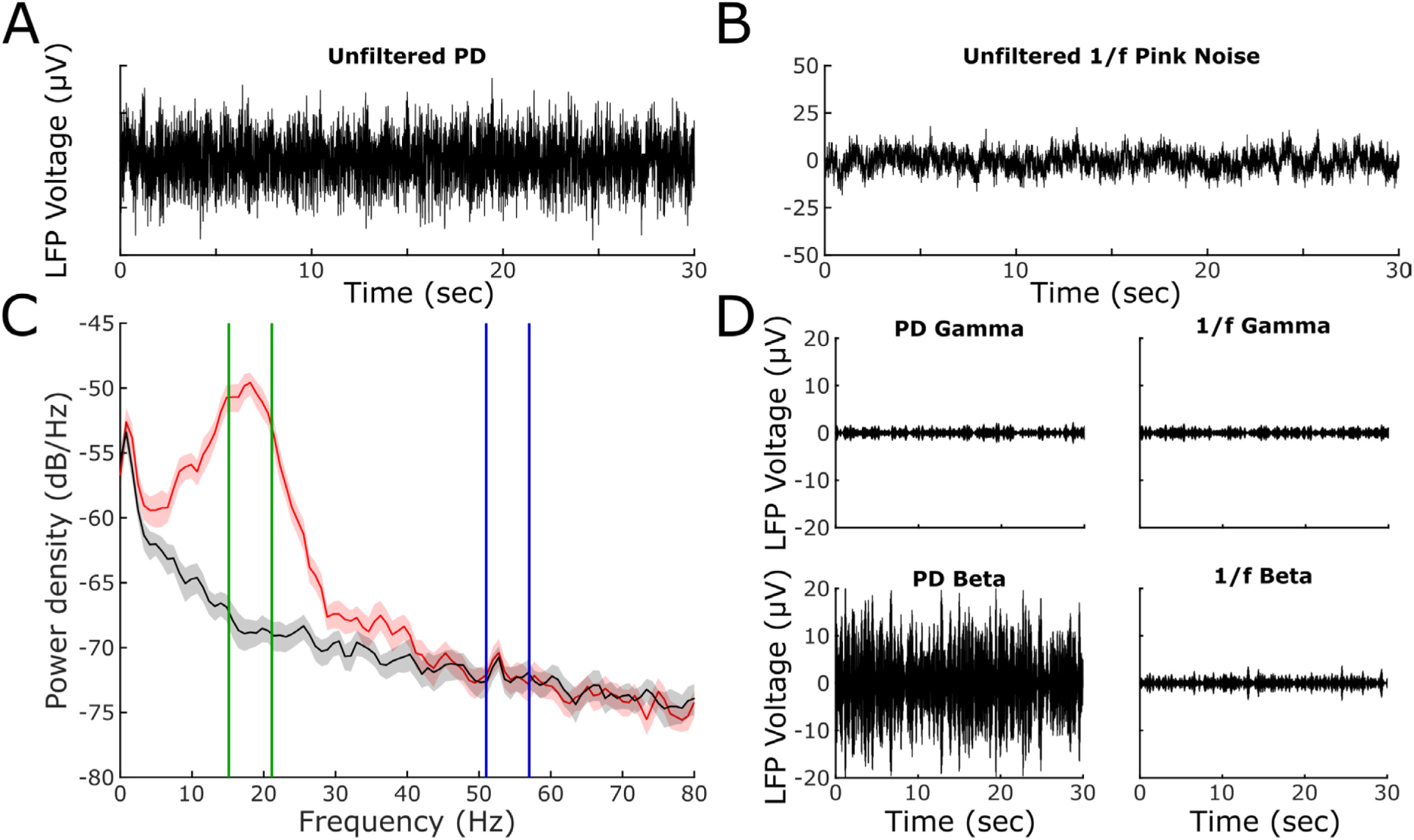
Local field potentials from a Parkinsonian patient are compared to simulated 1/f pink noise matched to the same roll off. Broad spectrum Parkinsonian local field potentials (A) are larger in amplitude than the simulated pink noise (B). Both signals are plotted on the same PSD (C), the power in the beta (13-30Hz) frequency range is elevated when compared with the gamma frequency range (40-70Hz). (D) Sub bands of the Parkinson’s and pink noise gamma (blue lines) and beta (green lines) from C are band pass filtered showing the elevation of power in the beta frequency range (Bottom Right).

When both signals were plotted on the same PSD, Fig. 1C, the power in a low gamma frequency band (40 - 70 Hz) overlapped the simulated physiological 1/f spectrum, whereas the power in the alpha and beta (8 - 30 Hz) frequency range was elevated above the 1/f spectrum. The LFPs from the 1/f and PD signals were each filtered in two 6 Hz bands: one in the band where the two PSDs overlapped (51 – 57 Hz, blue lines, Fig. 1C) and the other around the beta peak of the PSD spectrum (15.13 – 21.13 Hz, green vertical lines, Fig. 1C). In the overlapping band the PD and 1/f filtered signals were similar in amplitude, Fig. 1D top panels, however, the PD signal had greater amplitude than that in the 1/f signal in the elevated beta sub-band, Fig. 1D lower panels.

#### 3.2 Burst durations were similar in the PD and 1/f overlapping bands, establishing a physiological baseline from the PD spectrum

Figure 2A and B demonstrate the bandpass filtered signal from the overlapping band of the PD and 1/f spectrum, respectively. The bandpass filtered signals were squared and the envelope of those signals are shown in grey in Figure 2C and D respectively.

**Figure 2:**
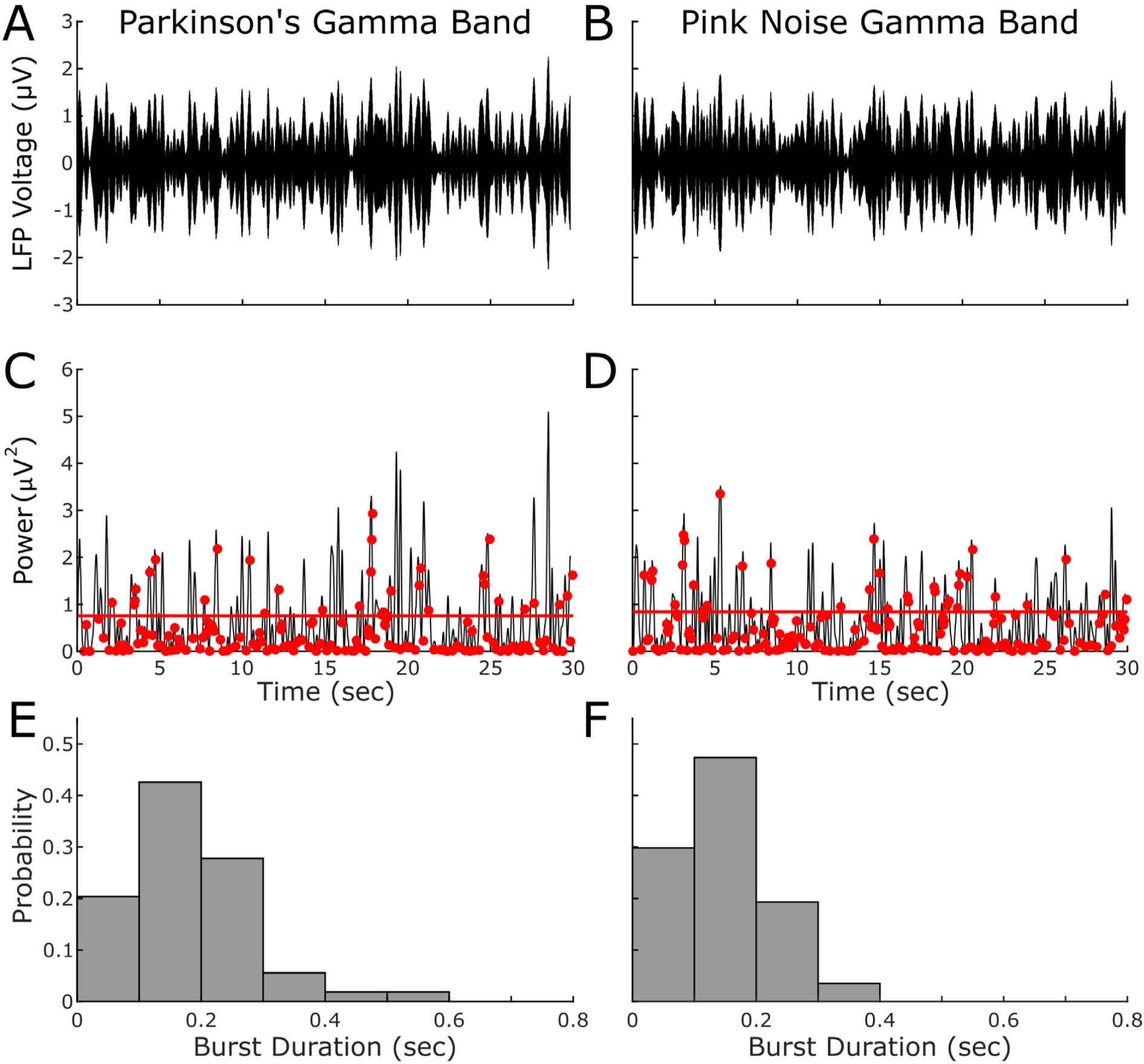
Bandpass filtered PD LFP and pink noise filtered for the gamma frequency band (CF = 54 Hz, BW = 6Hz). Parkinson’s gamma band data (A) and simulated pink noise data (B) are shown using the same axes. LFP power (black), identified troughs (red dots), thresholds (red line) and identified bursts using the Anderson method in PD (C) and pink noise (D). Histograms of the associated burst durations for PD (E) and pink noise (F) are similar.

A trough detection algorithm identified the local minima of the envelopes of the filtered squared signals (red dots, Figs. 2C and D), and the median power of these troughs was calculated for each signal. The red line in Figs. 2C and D represents 4 times the median power of the troughs and formed the baseline or threshold from which burst durations and power were calculated for each signal. The duration of a burst of power was defined as the period between ‘zero’ crossings of the envelope across the baseline; the mean power of the burst was the average power of the filtered and squared LFP signal bounded by the start and end time of the burst. The histograms in Fig. 2E and F demonstrate that the distributions and respective means of burst durations were similar between the overlapping PD and 1/f bands. The mean burst durations were 181 ± 102 ms and 153 ± 76 ms, respectively, t(109)=1.68; p=0.096, which were similar to those reported for physiological neural activity in non-human primates (40 – 120 ms, Feingold 2015). This supported the hypothesis that this portion of the PD spectrum comprised largely short duration (physiological) bursts of neural activity; we therefore designated the baseline of the PD overlapping band as the ‘physiological’ baseline that would be used to calculate burst dynamics in pathological or elevated bands of the PD spectrum.

#### 3.3 Calculation of burst durations depend on the choice of center frequency and bandwidth

Figure 3 demonstrates how varying filter bandwidth and center frequency affect burst duration. Mean burst durations of the 1000 episodes of pink noise were calculated with the signal filtered in successive bandwidth ranges from 1 – 10 Hz at each of the center frequencies of 10 Hz, 15 Hz, and 20 Hz.

**Figure 3:**
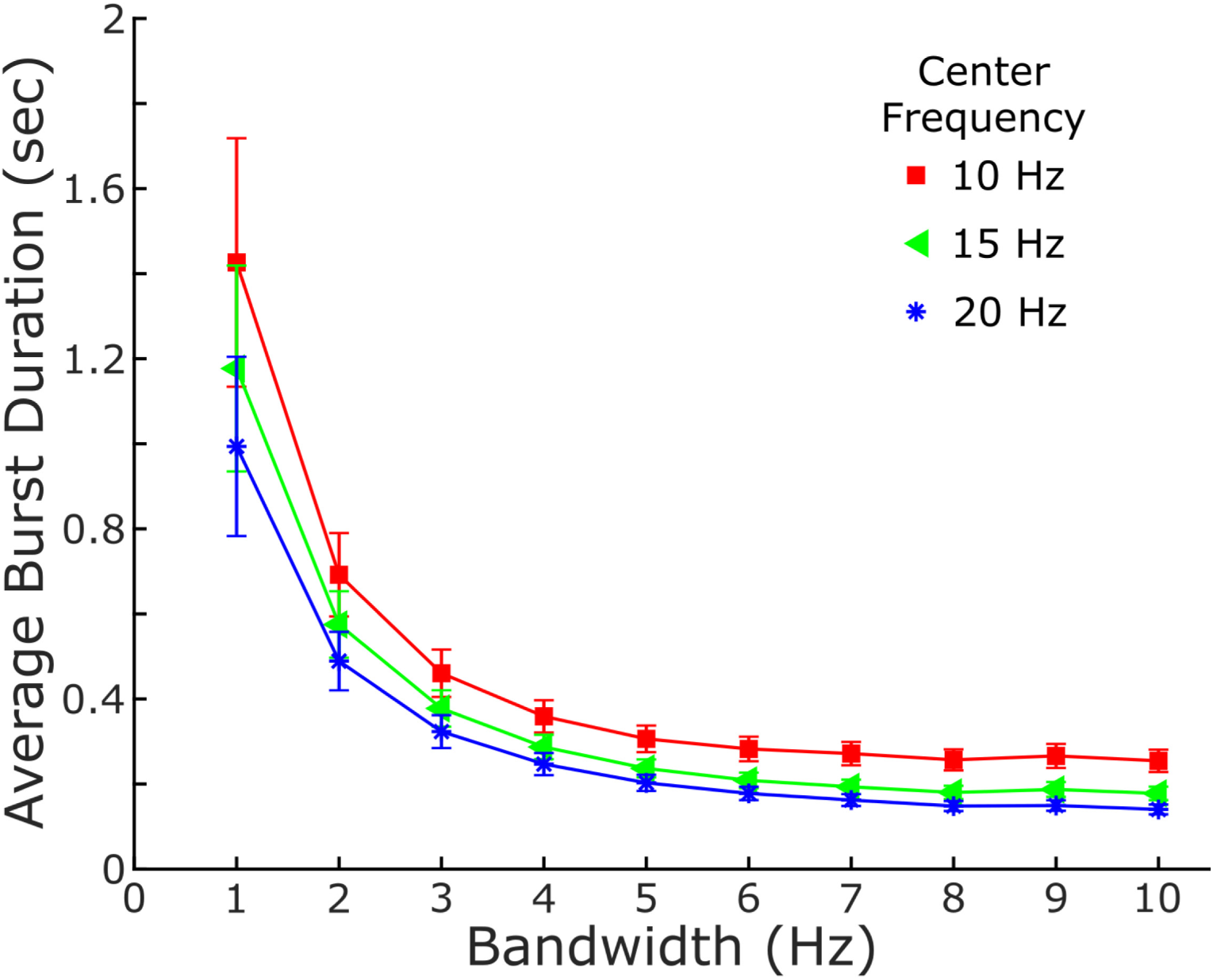
Burst durations are calculated for the Anderson method using 1000 runs of simulated pink noise. The effects of bandwidth and center frequency (10, 15, and 20 Hz) are shown. Error bars represent standard deviations of the mean.

The mean burst duration of the 1/f spectrum depended on the bandwidth of the filter used: it increased exponentially for narrow bandwidths and followed an asymptote towards little change at bandwidths longer than 6 Hz. At bandwidths greater than 2 Hz, mean burst durations of pink noise were higher at a center frequency of 10 Hz compared to 20 Hz. At bandwidths greater than 6 Hz, the mean burst durations were greater at the 10 Hz center frequency than at both 15 Hz and 20 Hz.

#### 3.4 The Anderson method captured a broader distribution of burst durations compared to methods using a threshold of power

Figure 4 demonstrates the differences in beta burst duration distribution when calculated using the Anderson method (Fig. 4B, E, H), the Feingold method (Feingold et al., 2015) (Fig. 4C, F, I), and the Tinkhauser method (Tinkhauser et al., 2017a) (Fig. 4D, G, J). The same thirty seconds of a PD LFP signal was used for all the methods and was filtered using a zero-phase 8th order Butterworth bandpass filter with a 6Hz bandwidth around the same center frequency (18.1 Hz); the filtered signal is shown in Fig. 4A.

**Figure 4:**
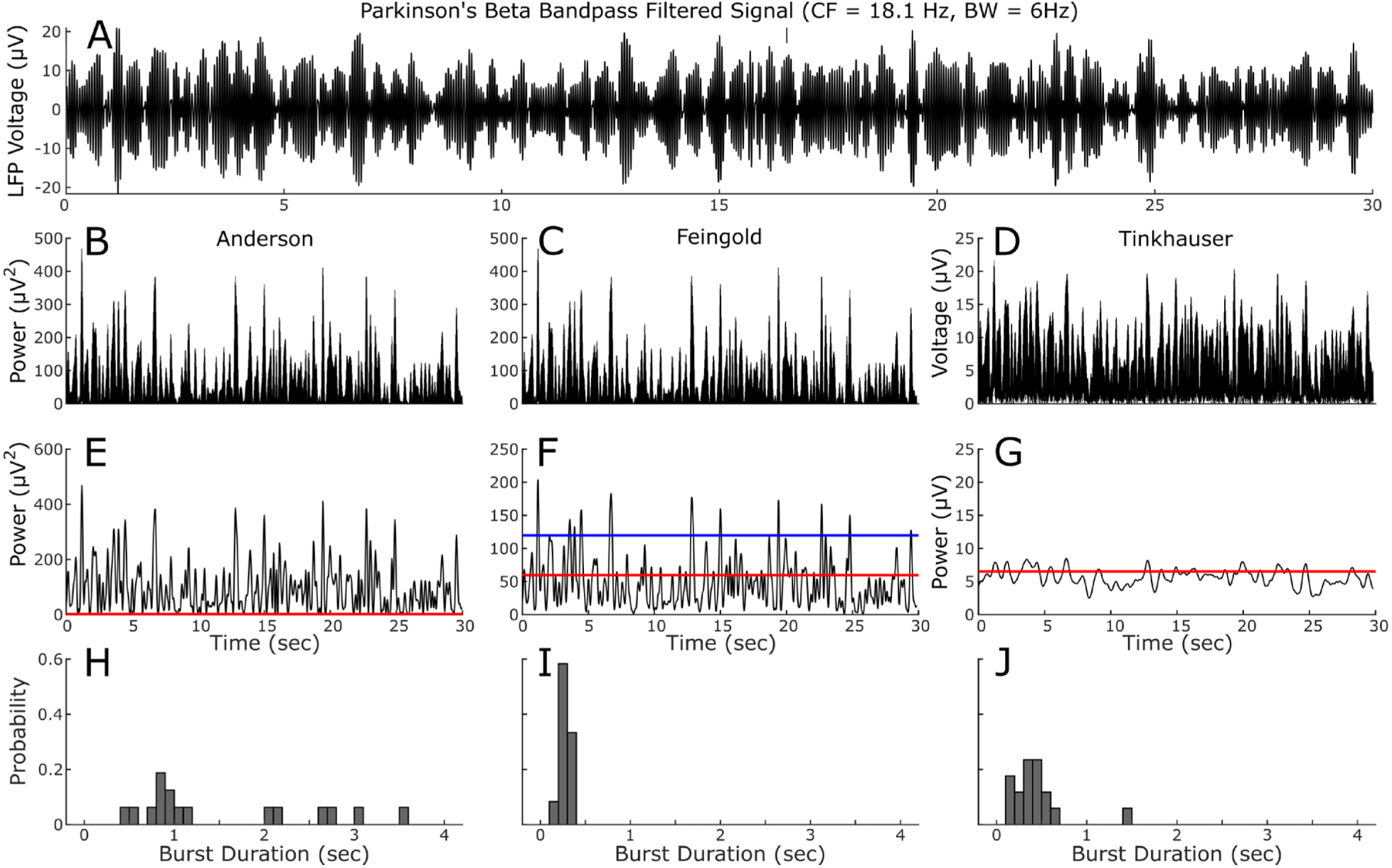
Comparison of the three burst duration methods (Anderson, Feingold, Tinkhauser) processed with the same center frequency (18.1Hz) and bandwidth (6 Hz). (A) Bandpass filtered LFP used for each analysis. Squared LFP and power envelope for the Anderson (B, E), Feingold (C, F), and Tinkhauser (D, G) methods. Histograms of the resulting burst durations for the Anderson (H), Feingold (I) and Tinkhauser (J) methods.

For the Anderson method, the bandpass filtered signal was squared (Fig. 4B), and an amplitude envelope was created by linearly connecting consecutive peaks of the filtered and squared LFP signal to form an envelope of the maximum power, (Fig. 4E). A baseline power was calculated by averaging the median trough amplitudes from 5 consecutive overlapping 6 Hz bands in the 45-63 Hz PD gamma spectrum, similar to that shown in Figure 2, from which fluctuations of power in the band in the PD spectrum would be measured. For the Feingold method the same bandpass filtered signal was squared, Fig.4C, and then smoothed using a Hanning window moving average filter, Fig. 4F. The Feingold method identified a burst when the signal power rose above a threshold that was 3 times the median power of the signal, blue line Fig. 4F, and used a threshold of 1.5 times the median power to calculate the burst durations, red line Fig. 4F (Feingold et al., 2015). The Tinkhauser method rectified the signal but did not square it, Fig. 4D (Tinkhauser et al., 2017a); it then smoothed the rectified signal using a 400 ms rectangular moving average filter, Fig. 4G, and identified and classified a burst using a power threshold, which could range from the 55^th^ to the 95^th^ percentile of the power in the smoothed signal (Tinkhauser, 75^th^ percentile shown). Signals above the red line were a burst while signals below the line were not a burst.

In all methods, the duration of a burst was calculated as the interval between successive crossings of the signal over the chosen baseline or threshold. Fig. 4H, I, J demonstrate how the distributions of burst durations from the same filtered signal differed, using the Anderson, Feingold and Tinkhauser methods respectively. The Anderson method resulted in the broadest distribution of beta burst durations (mean 1545 ± 1001 ms, range 405-3547 ms), the Feingold method had the narrowest distribution (mean 264 ± 65 ms, range 192-386 ms), and the Tinkhauser method had a distribution between the other methods (mean 439 ± 292 ms, range 147-1431 ms). The Anderson method differed in its definition of a burst from the other two methods, specifically in the intent to measure *all* fluctuations of power over a baseline rather than only the durations of higher power fluctuations using an event detection threshold, that was a percentage of the overall power of the spectrum.

#### 3.5 Resting state beta burst durations decrease during STN DBS at increasing intensity

The Anderson method was used to investigate the effect of increasing the intensity of STN DBS on resting state beta band burst durations, Figure 5.

**Figure 5:**
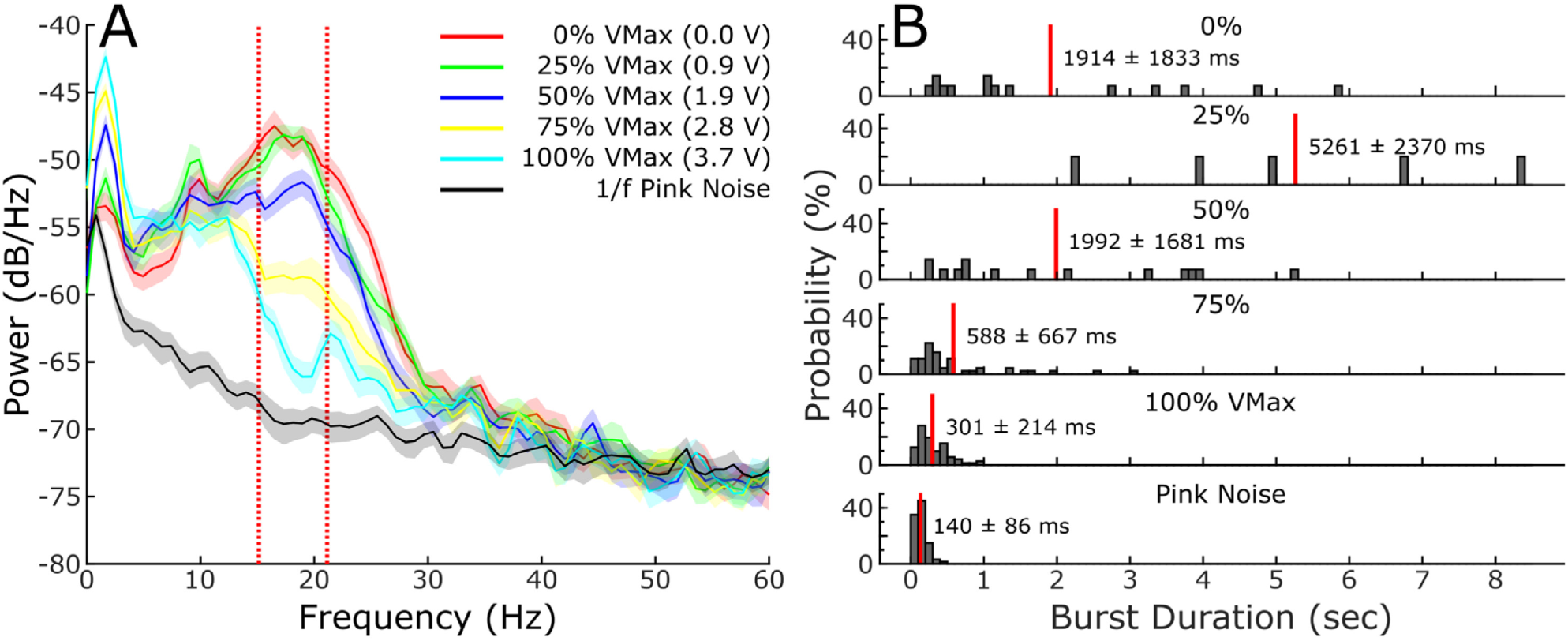
Beta power and burst durations are suppressed by deep brain stimulation. (A) Five different stimulation conditions (0%, 25%, 50%, 75% and 100% of clinical equivalent stimulation) are compared with simulated 1/f pink noise. Vertical red bars, a subset of the beta frequency band used for analysis in B that is highly dependent on stimulation intensity. (B) Burst durations calculated for the same 30 second windows are decreased with increasing levels of stimulation and are driven towards the normal burst duration expected with simulated pink noise. Mean burst durations are shown by labeled red bars.

Figure 5A demonstrates the effect of randomized epochs of STN DBS at different intensities on a representative resting state PD PSD; the pink noise estimate of the physiological 1/f PSD is demonstrated in black. The PD PSDs demonstrate attenuation of the resting state beta band power during STN DBS at increasing intensities (voltage) from 0% (no DBS, red PSD, Fig. 5A) to 100% of Vmax (3.7 V, light blue PSD, Fig. 5A). Burst durations calculated using the Anderson method, from a 6 Hz band centered on the peak beta band frequency (red vertical lines, Fig. 5B) revealed a broad distribution of burst durations in the resting state with no (0%) DBS, with a mean burst duration of 1914 ± 1833 ms. As the DBS intensity increased above 50% of Vmax, the distribution of beta band burst durations shifted towards shorter burst durations. During DBS at Vmax, the mean burst duration decreased to 301 ± 214 ms and the distribution was closer to that of pink noise (140 ± 86 ms). Across 9 PD subjects (16 STNs), mean burst duration at 0% (no) DBS was 933 ± 990 ms. Mean burst duration at Vmax was 250.6 ± 74.5 ms and was significantly shorter than the 0% DBS condition (t(15)=2.91; p=0.011).

## 4. Discussion

In this study, we introduce the Anderson method for measuring LFP power fluctuations (bursts) using a baseline rather than a threshold percentage of spectral power. The baseline was calculated from the troughs of power of a frequency band within the same PD STN spectrum, which overlapped the simulated physiological 1/f spectrum, and which had similar burst dynamics to physiological neural activity. The baseline was then used to calculate the burst durations in the elevated beta band of interest. Factors that influence burst duration and power calculation include the bandwidth and center frequency of the bandpass filter, the method of rectification (absolute value or squaring of the signal) and the type of smoothing applied to the signal. The Anderson method was used to investigate the effect of increasing intensities of STN DBS on resting state beta band burst durations and demonstrated that there was a significant reduction in mean beta band burst duration at the clinically equivalent DBS intensity compared to ‘OFF’ DBS. The distribution of burst durations shifted towards that of the simulated physiological signal as the DBS intensity increased. This suggests that the effect of DBS is to restore neural activity towards that of a physiological state, where beta band oscillations exist but are of shorter duration, closer to those demonstrated in the normal or non-Parkinsonian state (Deffains et al., 2018; Feingold et al., 2015; Murthy and Fetz, 1996, 1992).

### 4.1 Measurements of the dynamics of pathological beta band neural activity in Parkinson’s disease

Local field potentials (LFPs) represent the summed electrical activity of local neuronal networks and are classically represented by power spectral density (PSD) analysis and spectrograms, which do not provide adequate temporal and frequency resolution to accurately measure the temporal dynamics of neural activity. Several new methods for performing spectral deconstruction have emerged that utilize wavelets or bandpass filtering of local field potentials and these methods help to quantify changes in spectral power over time in narrower frequency bands, which has improved the ability to quantify the temporal nature of neuronal oscillatory activity. Currently, the published methods to measure the dynamics of beta band oscillatory activity have focused on event detection to define high power bursts. Aspects of the signal were only included in the burst analysis if and when instantaneous power rose above a threshold of the power of the LFP signal within that band or within the overall beta band, with or without a temporal threshold (e.g. power elevated for greater than a defined number of oscillatory cycles), (Cagnan et al., 2019; Deffains et al., 2018; Feingold et al., 2015; Lofredi et al., 2019b; Meidahl et al., 2019; Tinkhauser et al., 2017a, 2017b, 2018; Torrecillos et al., 2018). Most publications have followed the method described by Tinkhauser et al., who defined a burst when power exceeded a threshold of 75% of the power of the rectified smoothed signal. Activity below this power was discarded. In contrast, the Anderson method used a baseline power determined from a band not elevated above the 1/f signal, and whose burst dynamics resembled that of physiological neural activity. In this way, we sought to include as much of the signal from the band of interest as possible, including lower power fluctuations or bursts. This resulted in a broader range of burst durations compared to either the Tinkhauser or Feingold methods. The comparison of the Anderson and Feingold methods was more straightforward, as the signal was not as highly smoothed as that of Tinkhauser. The higher threshold in the Feingold method resulted in detection of high power, shorter duration bursts, whereas the lower power baseline of the Anderson method resulted in the inclusion of longer duration fluctuations, as well as shorter duration, lower power bursts. The emphasis on duration rather than power of beta bursts has been supported by Deffains and colleagues (Deffains et al., 2018), who demonstrated that STN beta burst duration was a more reliable metric than power to discriminate healthy and parkinsonian neural activity in non-human primates.

### 4.2 Burst duration varied with the choice of bandwidth, center frequency, and smoothing filters

Evaluation of 1000 samples of simulated 1/f neural activity filtered using different bandwidths at three different beta band center frequencies revealed that the mean burst duration had an inverse exponential relationship with bandwidth: at bandwidths shorter than 5 Hz, the mean burst duration progressively increased. Mean burst durations were shorter if the center frequency was 20 Hz compared to 10 Hz. This suggests that burst analysis methods should attempt to use the same bandwidth across each frequency band and take into account different center frequencies if applicable. For instance, the Feingold report used a large bandwidth (13-30 Hz or 17 Hz bandwidth) for all their recordings and a center frequency of ∼22 Hz, both of which would result in shorter burst durations of the 1/f signal, Fig. 3. In the original Tinkhauser analysis, the LFP signal was bandpass filtered in a 6 or 7 Hz bandwidth in eleven out of sixteen STNs and 3, 4, 5, or 13 Hz in the remaining STNs. The center frequencies varied from 16 – 31 Hz. This may have resulted in a small variation of burst duration based on different bandwidths and center frequencies chosen. The method used to rectify and/or smooth the raw LFP signal will also affect the burst duration and power.

### 4.3 Beta Burst duration as a biomarker of the efficacy of therapy and of Parkinsonian motor and gait impairment

In this study, we demonstrated that the mean resting state beta band burst duration and burst duration distributions progressively decreased with increasing intensities of DBS in 16 STNs. Prolonged resting state beta burst durations have been correlated with increasing PD motor severity (Tinkhauser et al., 2017a) and prolonged beta burst durations during gait have been shown to be a neural feature of the freezer phenotype and freezing of gait (FOG) (Anidi et al., 2018). Mean resting state beta burst durations were shorter on compared to off medication (Lofredi et al., 2019a; Tinkhauser et al., 2017b) and both 60 Hz and 140 Hz STN DBS decreased mean beta burst duration in freezers, and improved FOG. Only one study has suggested that burst durations may not be altered during DBS (Schmidt et al., 2020).

This growing body of evidence supports the hypothesis that longer duration beta bursts in subcortical structures, equating to longer periods of beta oscillations and synchrony, represent pathological or impaired sensorimotor processing in Parkinson’s disease (Anidi et al., 2018; Cagnan et al., 2019; Deffains et al., 2018; Lofredi et al., 2019b; Tinkhauser et al., 2018, 2017a, 2017b) and that one mechanism of STN DBS is to remove pathological sensorimotor network processing, while leaving intact normal physiological processing (Anidi et al., 2018; Holt et al., 2019; Tinkhauser et al., 2017a; Torrecillos et al., 2018). The discovery that averaged beta band power was attenuated during increasing intensities of DBS led to its successful use as the control variable in closed loop DBS studies (Afzal et al., 2019; Eusebio et al., 2011; Little et al., 2016a, 2016b, 2013; Piña-Fuentes et al., 2019, 2017; Rosa et al., 2017, 2015; Syrkin-Nikolau et al., 2017; Velisar et al., 2019; Whitmer et al., 2012). Similarly, the accumulating evidence suggests that beta burst duration will be a functionally relevant, patient specific control variable for closed loop DBS.

## 5. Conclusions

The Anderson method is a novel method for characterizing bursts of activity in local field potentials and can be applied to any band of interest in the LFP spectrum. We validated this method using pathological activity from people with Parkinson’s disease and simulated physiological neural activity. The baseline was determined from an “inert” band within the PD spectrum, which overlapped with the simulated 1/f physiological spectrum and had similar burst duration distributions and mean burst durations. Burst durations calculated using the Anderson method captured a wide range of Parkinsonian spectral activity: it revealed mainly short burst durations in the inert band and captured the longest and broadest distribution of burst durations in a pathological beta band compared to other methods. The Anderson method demonstrated that increased intensities of subthalamic neurostimulation shortened mean beta band burst durations and shifted their distribution towards that of the physiological spectrum. We suggest that this novel method is well suited to quantify the full range of fluctuations in beta band neural activity in the PD brain and is applicable to other bands of interest for other neuropsychological diseases. Beta band burst duration is a functionally relevant, patient specific control variable that can be used for closed loop neuromodulation algorithms in Parkinson’s disease.

## Author contributions

**Ross Anderson:** Conceptualization, Methodology, Software, Validation, Formal analysis, Investigations, Data Curation, Writing – Original Draft & Review & Editing, Visualization, Supervision, Funding acquisition. **Raumin Neuville:** Conceptualization, Validation, Formal analysis, Investigation, Writing – Review & Editing. **Yasmine Kehnemouyi:** Formal analysis, Investigation, Writing – Original Draft & Review and Editing. **Chioma Anidi:** Methodology, Validation, Formal Analysis. **Matthew Petrucci:** Methodology, Validation, Investigation, Supervision. **Jordan Parker:** Validation, Investigation. **Anca Velisar:** Software, Investigation. **Helen Bronte-Stewart:** Conceptualization, Methodology, Writing – Original Draft & Review & Editing, Supervision, Funding acquisition.

## Acknowledgements

We would like to thank the members of the Human Motor Control and Neuromodulation Lab and the participants in the study who without their help, none of this would be possible.

## References

Afzal, M.F., Velisar, A., Anidi, C., Neuville, R., Prabhakar, V., Bronte-Stewart, H., 2019. Proceedings #61: Subthalamic Neural Closed-loop Deep Brain Stimulation for Bradykinesia in Parkinson’s Disease. Brain Stimulation 12, e152.#x2013;e154. https://doi.org/10.1016/j.brs.2019.03.019

Anidi, C., O’Day, J.J., Anderson, R.W., Afzal, M.F., Syrkin-Nikolau, J., Velisar, A., Bronte-Stewart, H.M., 2018. Neuromodulation targets pathological not physiological beta bursts during gait in Parkinson’s disease. Neurobiol. Dis. 120, 107–117. https://doi.org/10.1016/j.nbd.2018.09.004

Bergman, H., Wichmann, T., Karmon, B., DeLong, M.R., 1994. The primate subthalamic nucleus. II. Neuronal activity in the MPTP model of parkinsonism. J. Neurophysiol. 72, 507–520. https://doi.org/10.1152/jn.1994.72.2.507

Brontë-Stewart, H., Louie, S., Batya, S., Henderson, J.M., 2010. Clinical motor outcome of bilateral subthalamic nucleus deep-brain stimulation for Parkinson’s disease using imageguided frameless stereotaxy. Neurosurgery 67, 1088–1093; discussion 1093. https://doi.org/10.1227/NEU.0b013e3181ecc887

Brown, P., 2003. Oscillatory nature of human basal ganglia activity: relationship to the pathophysiology of Parkinson’s disease. Mov. Disord. 18, 357–363. https://doi.org/10.1002/mds.10358

Cagnan, H., Mallet, N., Moll, C.K.E., Gulberti, A., Holt, A.B., Westphal, M., Gerloff, C., Engel, A.K., Hamel, W., Magill, P.J., Brown, P., Sharott, A., 2019. Temporal evolution of beta bursts in the parkinsonian cortical and basal ganglia network. Proc. Natl. Acad. Sci. U.S.A. 116, 16095–16104. https://doi.org/10.1073/pnas.1819975116

Courtemanche, R., Fujii, N., Graybiel, A.M., 2003. Synchronous, focally modulated beta-band oscillations characterize local field potential activity in the striatum of awake behaving monkeys. J. Neurosci. 23, 11741–11752.

Deffains, M., Iskhakova, L., Katabi, S., Israel, Z., Bergman, H., 2018. Longer β oscillatory episodes reliably identify pathological subthalamic activity in Parkinsonism. Mov. Disord. 33, 1609–1618. https://doi.org/10.1002/mds.27418

Eusebio, A., Thevathasan, W., Doyle Gaynor, L., Pogosyan, A., Bye, E., Foltynie, T., Zrinzo, L., Ashkan, K., Aziz, T., Brown, P., 2011. Deep brain stimulation can suppress pathological synchronisation in parkinsonian patients. J. Neurol. Neurosurg. Psychiatry 82, 569–573. https://doi.org/10.1136/jnnp.2010.217489

Feingold, J., Gibson, D.J., DePasquale, B., Graybiel, A.M., 2015. Bursts of beta oscillation differentiate postperformance activity in the striatum and motor cortex of monkeys performing movement tasks. PNAS 112, 13687–13692. https://doi.org/10.1073/pnas.1517629112

Hammond, C., Bergman, H., Brown, P., 2007. Pathological synchronization in Parkinson’s disease: networks, models and treatments. Trends Neurosci. 30, 357–364. https://doi.org/10.1016/j.tins.2007.05.004

He, B.J., 2014. Scale-free brain activity: past, present, and future. Trends Cogn. Sci. (Regul. Ed.) 18, 480–487. https://doi.org/10.1016/j.tics.2014.04.003

Holt, A.B., Kormann, E., Gulberti, A., Pötter-Nerger, M., McNamara, C.G., Cagnan, H., Baaske, M.K., Little, S., Köppen, J.A., Buhmann, C., Westphal, M., Gerloff, C., Engel, A.K., Brown, P., Hamel, W., Moll, C.K.E., Sharott, A., 2019. Phase-Dependent Suppression of Beta Oscillations in Parkinson’s Disease Patients. J. Neurosci. 39, 1119–1134. https://doi.org/10.1523/JNEUROSCI.1913-18.2018

Kühn, A.A., Kempf, F., Brücke, C., Gaynor Doyle, L., Martinez-Torres, I., Pogosyan, A., Trottenberg, T., Kupsch, A., Schneider, G.-H., Hariz, M.I., Vandenberghe, W., Nuttin, B., Brown, P., 2008. High-frequency stimulation of the subthalamic nucleus suppresses oscillatory beta activity in patients with Parkinson’s disease in parallel with improvement in motor performance. J. Neurosci. 28, 6165–6173. https://doi.org/10.1523/JNEUROSCI.0282-08.2008

Kühn, A.A., Kupsch, A., Schneider, G.-H., Brown, P., 2006. Reduction in subthalamic 8-35 Hz oscillatory activity correlates with clinical improvement in Parkinson’s disease: STN activity and motor improvement. European Journal of Neuroscience 23, 1956–1960. https://doi.org/10.1111/j.1460-9568.2006.04717.x

Little, S., Beudel, M., Zrinzo, L., Foltynie, T., Limousin, P., Hariz, M., Neal, S., Cheeran, B., Cagnan, H., Gratwicke, J., Aziz, T.Z., Pogosyan, A., Brown, P., 2016a. Bilateral adaptive deep brain stimulation is effective in Parkinson’s disease. Journal of Neurology, Neurosurgery & Psychiatry 87, 717–721. https://doi.org/10.1136/jnnp-2015-310972

Little, S., Pogosyan, A., Neal, S., Zavala, B., Zrinzo, L., Hariz, M., Foltynie, T., Limousin, P., Ashkan, K., FitzGerald, J., Green, A.L., Aziz, T.Z., Brown, P., 2013. Adaptive deep brain stimulation in advanced Parkinson disease. Annals of Neurology 74, 449–457. https://doi.org/10.1002/ana.23951

Little, S., Tripoliti, E., Beudel, M., Pogosyan, A., Cagnan, H., Herz, D., Bestmann, S., Aziz, T., Cheeran, B., Zrinzo, L., Hariz, M., Hyam, J., Limousin, P., Foltynie, T., Brown, P., 2016b. Adaptive deep brain stimulation for Parkinson’s disease demonstrates reduced speech side effects compared to conventional stimulation in the acute setting. Journal of Neurology, Neurosurgery & Psychiatry jnnp-2016-313518. https://doi.org/10.1136/jnnp-2016-313518

Lofredi, R., Neumann, W.-J., Brücke, C., Huebl, J., Krauss, J.K., Schneider, G.-H., Kühn, A.A., 2019a. Pallidal beta bursts in Parkinson’s disease and dystonia. Mov. Disord. 34, 420–424. https://doi.org/10.1002/mds.27524

Lofredi, R., Tan, H., Neumann, W.-J., Yeh, C.-H., Schneider, G.-H., Kühn, A.A., Brown, P., 2019b. Beta bursts during continuous movements accompany the velocity decrement in Parkinson’s disease patients. Neurobiol. Dis. 127, 462–471. https://doi.org/10.1016/j.nbd.2019.03.013

Meidahl, A.C., Moll, C.K.E., van Wijk, B.C.M., Gulberti, A., Tinkhauser, G., Westphal, M., Engel, A.K., Hamel, W., Brown, P., Sharott, A., 2019. Synchronised spiking activity underlies phase amplitude coupling in the subthalamic nucleus of Parkinson’s disease patients. Neurobiol. Dis. 127, 101–113. https://doi.org/10.1016/j.nbd.2019.02.005

Murthy, V.N., Fetz, E.E., 1996. Synchronization of neurons during local field potential oscillations in sensorimotor cortex of awake monkeys. J. Neurophysiol. 76, 3968–3982.

Murthy, V.N., Fetz, E.E., 1992. Coherent 25-to 35-Hz oscillations in the sensorimotor cortex of awake behaving monkeys. Proc. Natl. Acad. Sci. U.S.A. 89, 5670–5674.

Piña-Fuentes, D., Beudel, M., Little, S., Brown, P., Oterdoom, D.L.M., van Dijk, J.M.C., 2019. Adaptive deep brain stimulation as advanced Parkinson’s disease treatment (ADAPT study): protocol for a pseudo-randomised clinical study. BMJ Open 9, e029652. https://doi.org/10.1136/bmjopen-2019-029652

Piña-Fuentes, D., Little, S., Oterdoom, M., Neal, S., Pogosyan, A., Tijssen, M.A.J., van Laar, T., Brown, P., van Dijk, J.M.C., Beudel, M., 2017. Adaptive DBS in a Parkinson’s patient with chronically implanted DBS: A proof of principle. Mov. Disord. 32, 1253–1254. https://doi.org/10.1002/mds.26959

Quinn, E.J., Blumenfeld, Z., Velisar, A., Koop, M.M., Shreve, L.A., Trager, M.H., Hill, B.C., Kilbane, C., Henderson, J.M., Brontë-Stewart, H., 2015. Beta oscillations in freely moving Parkinson’s subjects are attenuated during deep brain stimulation: Beta Oscillations In Free Moving PD Subjects. Movement Disorders 30, 1750–1758. https://doi.org/10.1002/mds.26376

Rosa, M., Arlotti, M., Ardolino, G., Cogiamanian, F., Marceglia, S., Di Fonzo, A., Cortese, F., Rampini, P.M., Priori, A., 2015. Adaptive deep brain stimulation in a freely moving parkinsonian patient: Letters: New Observations. Movement Disorders 30, 1003–1005. https://doi.org/10.1002/mds.26241

Rosa, M., Arlotti, M., Marceglia, S., Cogiamanian, F., Ardolino, G., Fonzo, A.D., Lopiano, L., Scelzo, E., Merola, A., Locatelli, M., Rampini, P.M., Priori, A., 2017. Adaptive deep brain stimulation controls levodopa-induced side effects in Parkinsonian patients. Mov. Disord. 32, 628–629. https://doi.org/10.1002/mds.26953

Schmidt, S.L., Peters, J.J., Turner, D.A., Grill, W.M., 2020. Continuous deep brain stimulation of the subthalamic nucleus may not modulate beta bursts in patients with Parkinson’s disease. Brain Stimul 13, 433–443. https://doi.org/10.1016/j.brs.2019.12.008

Shreve, L.A., Velisar, A., Malekmohammadi, M., Koop, M.M., Trager, M., Quinn, E.J., Hill, B.C., Blumenfeld, Z., Kilbane, C., Mantovani, A., Henderson, J.M., Brontë-Stewart, H., 2017. Subthalamic oscillations and phase amplitude coupling are greater in the more affected hemisphere in Parkinson’s disease. Clin Neurophysiol 128, 128–137. https://doi.org/10.1016/j.clinph.2016.10.095

Syrkin-Nikolau, J., Koop, M.M., Prieto, T., Anidi, C., Afzal, M.F., Velisar, A., Blumenfeld, Z., Martin, T., Trager, M., Bronte-Stewart, H., 2017. Subthalamic neural entropy is a feature of freezing of gait in freely moving people with Parkinson’s disease. Neurobiol. Dis. 108, 288–297. https://doi.org/10.1016/j.nbd.2017.09.002

Tinkhauser, G., Pogosyan, A., Little, S., Beudel, M., Herz, D.M., Tan, H., Brown, P., 2017a. The modulatory effect of adaptive deep brain stimulation on beta bursts in Parkinson’s disease. Brain 140, 1053–1067. https://doi.org/10.1093/brain/awx010

Tinkhauser, G., Pogosyan, A., Tan, H., Herz, D.M., Kühn, A.A., Brown, P., 2017b. Beta burst dynamics in Parkinson’s disease OFF and ON dopaminergic medication. Brain. https://doi.org/10.1093/brain/awx252

Tinkhauser, G., Torrecillos, F., Duclos, Y., Tan, H., Pogosyan, A., Fischer, P., Carron, R., Welter, M.-L., Karachi, C., Vandenberghe, W., Nuttin, B., Witjas, T., Régis, J., Azulay, J.-P., Eusebio, A., Brown, P., 2018. Beta burst coupling across the motor circuit in Parkinson’s disease. Neurobiol. Dis. 117, 217–225. https://doi.org/10.1016/j.nbd.2018.06.007

Torrecillos, F., Tinkhauser, G., Fischer, P., Green, A.L., Aziz, T.Z., Foltynie, T., Limousin, P., Zrinzo, L., Ashkan, K., Brown, P., Tan, H., 2018. Modulation of Beta Bursts in the Subthalamic Nucleus Predicts Motor Performance. J. Neurosci. 38, 8905–8917. https://doi.org/10.1523/JNEUROSCI.1314-18.2018

Trager, M.H., Koop, M.M., Velisar, A., Blumenfeld, Z., Nikolau, J.S., Quinn, E.J., Martin, T., Bronte-Stewart, H., 2016. Subthalamic beta oscillations are attenuated after withdrawal of chronic high frequency neurostimulation in Parkinson’s disease. Neurobiology of Disease 96, 22–30. https://doi.org/10.1016/j.nbd.2016.08.003

Velisar, A., Syrkin-Nikolau, J., Blumenfeld, Z., Trager, M.H., Afzal, M.F., Prabhakar, V., Bronte-Stewart, H., 2019. Dual threshold neural closed loop deep brain stimulation in Parkinson disease patients. Brain Stimul 12, 868–876. https://doi.org/10.1016/j.brs.2019.02.020

Welch, P., 1967. The use of fast Fourier transform for the estimation of power spectra: A method based on time averaging over short, modified periodograms. IEEE Transactions on Audio and Electroacoustics 15, 70–73. https://doi.org/10.1109/TAU.1967.1161901

Whitmer, D., de Solages, C., Hill, B., Yu, H., Henderson, J.M., Bronte-Stewart, H., 2012. High frequency deep brain stimulation attenuates subthalamic and cortical rhythms in Parkinson’s disease. Front Hum Neurosci 6, 155. https://doi.org/10.3389/fnhum.2012.00155

